# UCsim2: 2D Structured Illumination Microscopy using UC2

**DOI:** 10.1101/2021.01.08.425840

**Authors:** Haoran Wang, René Lachmann, Barbora Marsikova, Rainer Heintzmann, Benedict Diederich

## Abstract

State-of-the-art microscopy techniques enable the imaging of sub-diffraction barrier biological structures at the price of high-costs or lacking transparency. We try to reduce some of these barriers by presenting a super-resolution upgrade to our recently presented open-source optical toolbox UC2. Our new injection moulded parts allow larger builds with higher precision. The 4× lower manufacturing tolerance compared to 3D printing makes assemblies more reproducible. By adding consumer-grade available open-source hardware such as digital mirror devices (DMD) and laser projectors we demonstrate a compact 3D multimodal setup that combines image scanning microscopy (ISM) and structured illumination microscopy (SIM). We demonstrate a gain in resolution and optical sectioning using the two different modes compared to the widefield limit by imaging Alexa Fluor 647- and SiR-stained HeLa cells. We compare different objective lenses and by sharing the designs and manuals of our setup, we make super-resolution imaging available to everyone.

## Background

During the last decade biotechnology has seen rapid development and became an important factor for economy. This is especially visible in the pandemic of the SARS-CoV-2 coronavirus, which has led reams of research groups to support the invention and development of tools for new insights into questions about the unknown virus[1],[2]. Optical methods, such as microscopy, have emerged as an important tool for biologists to understand the underlying mechanisms of, for example, virus entry or cell-cell interaction. The ability to localize a specifically labelled target at the nanometre scale enables correlative measurements that can give the scientist a more comprehensive picture of the sample under study. Traditional light microscopy methods are bound by the diffraction limit given by Ernst Abbe[3] and cannot resolve small structural details. However, novel techniques in which sophisticated optical setups, powerful reconstruction algorithms, and biochemical protocols work hand in hand can circumvent this limit and provide new insights into previously invisible worlds.

Yet, various methods for optical super-resolution, such as stimulated emission depletion (STED), image scanning microscopy (ISM[4], [5]), direct stochastic optical reconstruction microscopy (*d*STORM[6]) and structured illumination microscopy (SIM[7]), have one thing in common: a strong correlation between price, often accompanied by complexity of the setup, and achieved resolution. This leads to a separation between builders and users thereby harming interdisciplinary research. Biology workshops and interactive microscopy courses, such as those offered by the European Molecular Biology Organization (EMBO), try to combine both worlds through lectures and hands-on experience. However, commercial front-end microscopes remain largely a black box, not only because of their complexity but also because of their proprietary nature. This complicates the adaptation of innovative do-it-yourself (DIY) imaging solutions, as necessary information of the hardware equipment is often not provided by the manufacturers.

Open-source projects from software and hardware try to accelerate research and make it more reproducible by providing freely accessible interfaces, easy interaction and extension of existing projects. Projects such as the monolithic SMLM system miCube[8], the 3D printed OpenFlexure microscope[9] or the portable microscopy framework Squid[10] have proven to enable state-of-the-art microscopy at a reasonable price. Similarly, the DMD-based low-cost fairSIM realizes structured illumination microscopy in high safety biological labs by use of low-cost components in combination with customized software[11]. Likewise, a MEMS-based setup[12] creating an incoherently temporally varying excitation pattern could demonstrate a simple realisation of an ISM, which reduces complexity and costs by relying on commercially available laser video projectors. With the modularization and the resulting simplification of optical setups, we have recently been able to demonstrate with the open-source optical toolbox UC2[13] that sophisticated setups can also be accomplished outside an optical laboratory by educated users from other fields.

In this work we aim to continue the democratization of super-resolution tools and show how the UC2 toolbox can be used to realize methods such as ISM and SIM. This work consists of three parts. Firstly, we present the improved basic cube now enabling higher stability, setup-compatibility across different 3D-printers and fewer parts needed for building and stacking. Secondly, we demonstrate how two-dimensional ISM and SIM can be realized within the UC2 framework by using more complex module assemblies and adding opto-electronic components such as consumer laser video projectors or open-source digital mirror device (DMD) based displays. We apply these methods to biological samples and demonstrate the gain in optical sectioning and higher optical resolution as compared to a wide-field illumination scheme. The here created combination of ISM and SIM in one multimodal device has the smallest footprint and price of its kind to our knowledge and comes with a comprehensive documentation to allow people reproducing it.

## Methods

### Upgrading UC2 with Injection Moulded (IM) Parts

UC2 (literally “You. See. Too.”) is based on the Fourier optical principle, where Fourier- and image planes alternate and focal planes of lenses coincide. This simplifies development and building of optical systems, since modules with optical functionality can then just be stacked. The original UC2 system, which is described in detail in [13], relies entirely on 3D printed cubes that host a variety of different optical components with the help of customized inserts.

We demonstrated that the 3D printed cubes can be used to build multiple identical microscopes. This allows parallelizing experiments. 3D printing proved itself to be a valuable method for rapid prototyping and due to that it is possible to develop an imaging system for a given application in a short time. However, once a certain system is developed and optimised, 3D printing is no longer the method of choice. The printing and assembly time of the modules becomes an issue especially during upscaling towards multiple parallel setups. Furthermore, the designs presented in our previous publication[13] suffer from slight differences in scale between different 3D-printers with the stability and reproducibility depending also on the choice of the printing material.

Here, we introduce a novel injection-mouldable design, which does not require additional screws as compared to the previous version (v2) in Figure 1a). The new cube (v3, Figure 1b) now relies on a point-symmetric structure, where the two halves can simply be stuck together with a mechanical locking mechanism. The assembled cube finds its place on a 5 mm thick tiled baseplate, which can be extended infinitely in X/Y due to its puzzle mechanism (Figure 1c). Only two moulds are needed - one for the cube-half and one for the baseplate puzzle piece - thereby reducing costs. Using pins and fitting holes, the connection is maintained without any play over a long time, with a defined mechanical stop and offers a much higher precision in all directions. This mechanism automatically centres the part and avoids mechanical ambiguities. During the designing process, we ensured that the new version is downward-compatible with previously presented 3D-printable cubes[13]. By using M5 set screws in the positions of the casted pins, this novel positive locking mechanism can also be used with fully 3D-printed modules, even though this design is not optimized for printing.

**Figure 1 –.**
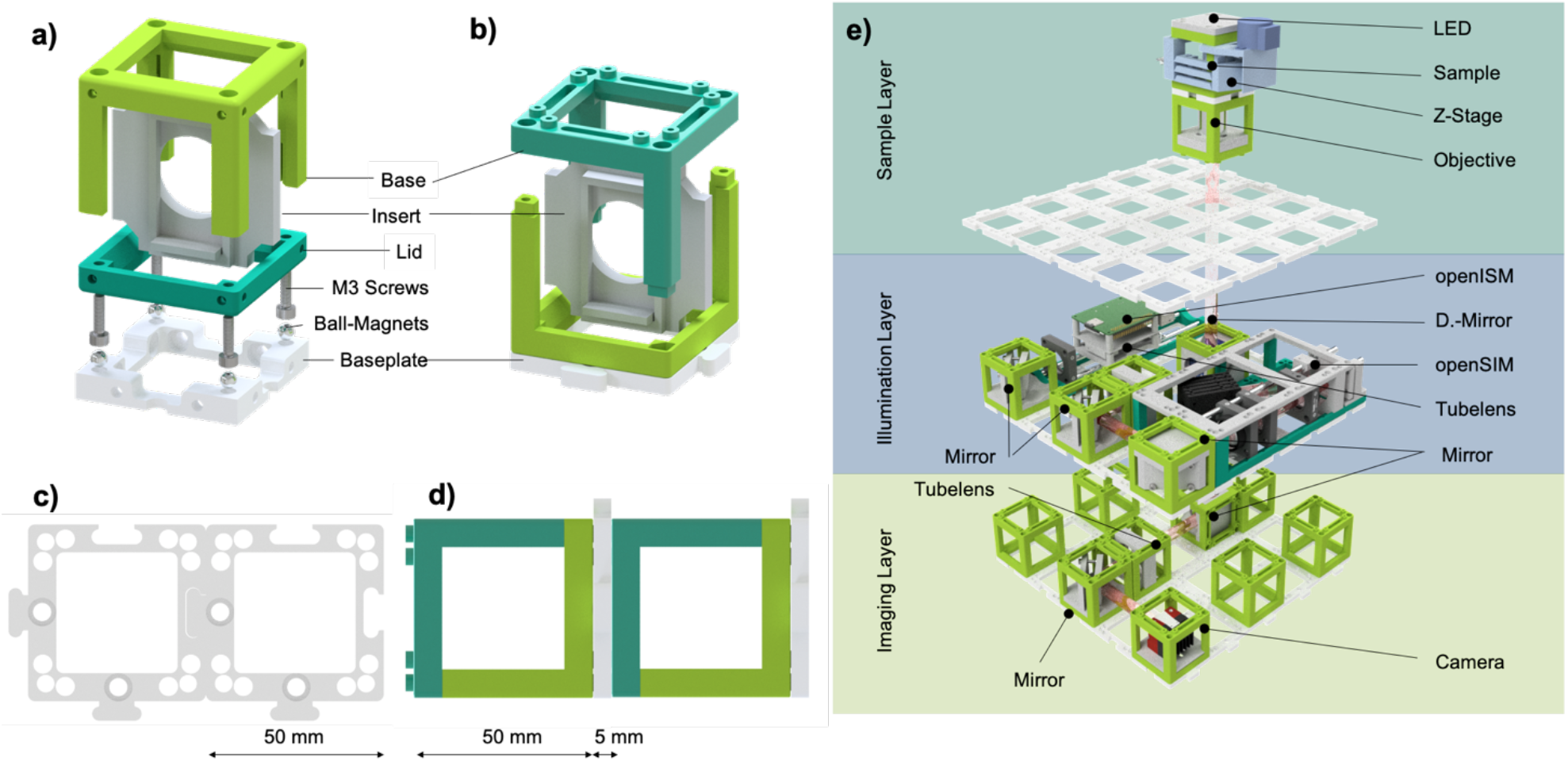
Injection Moulding Upgrade of the Optical Building Block: a) The previously presented cube assembly (V2) consists of a base and a lid screwed together using cylindrical M3 screws placed on ball magnets sitting in a baseplate. The insert is form locked inside the assembly. b) The new injection mould design consists of two point-symmetric bases held together by a form lock, omitting the use of any screws. The puzzle piece-based baseplate c) allows form-fit based connection of the cubes in horizontal direction d). e) Visualization of the 3D stacked assembly which combines ISM and SIM in one setup. The “Imaging Layer” holds two kinematic mirror mounts, tube-lens, (CMOS) camera and several empty cubes for stability. The “Illumination Layer” consists of the light engines for ISM (grid-pattern) and SIM (2-points). A dichromatic reflector passes the illumination-light to the back focal plane (BFP) of the objective and on reflection transmits the fluorescence to the imaging-layer. The Z-stage focuses the sample and incorporates an LED for standard brightfield imaging.

The new fit-mechanism allows for direct stacking along a 3^rd^ dimension by connecting baseplate-cube-pairs. This further increases mechanical stability since the cubes are fixed on both sides (Figure 1d). This “sandwich”-like design simplifies alignment of optical elements by dividing the setup in functional layers (e.g., imaging, illumination, and sample) and pre-aligning each layer before stacking them. The inherently induced 5 mm offset caused by the baseplate layer must be considered when designing UC2 setups (Figure 1e).

The design-files as well as the bill of material (BOM) and detailed instructions on how to build this module can be found in the online repository^1^,[14]. The overall component costs for the whole multimodal system including all optics, electronics, a camera and the computational device is in the range of 2000€ (see Table 1).

### Image Scanning Microscopy (ISM)

Classical confocal microscopy offers high optical sectioning by imaging a point source (e.g. a pinhole) into the sample plane and reimaging the illuminated spot onto a detector^15^. A second pinhole in front of the detector which is “con-focal” (i.e. the pinhole and illumination focus being at conjugated locations) to the illumination pinhole can block out-of-focus light. This process can be sped up with the help of a disc featuring multiple pinholes[15] or by rapidly scanning the illumination source (e.g. a laser) laterally over the sample plane, effectively writing the illumination pattern onto the sample and blanking unwanted illumination positions, whilst detecting the scattered or fluorescent signal on an area detector (e.g. a camera) for each illumination pattern.

In ISM, the image is sampled with multiple detector pixels such that in-place[16] or post-processing[4] pixel reassignment leads to higher resolution and a more efficient photon-usage[16].

Compared imaging setups equipped with a single detector, faster image acquisition can be achieved by multi-spot imaging thereby making use of multiple detectors or different regions of a camera at the same time[17]–[19]. Under these circumstances one acquired image 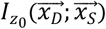 can be approximated in the noise-free limit as follows:

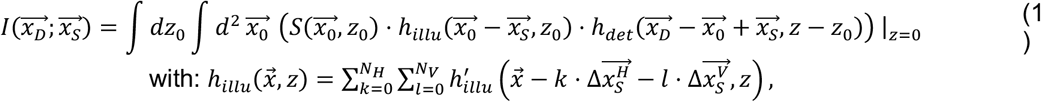

where *h*_*illu*_ is the total illumination pattern and 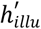 the illumination intensity point spread function (PSF) corresponding to a single focussed point, *h*_*det*_ the detection PSF, 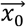 sample coordinates, 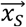 the scan position, 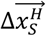 and 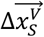 the vectors defining the illumination lattice, and 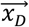 the detector coordinate system. Only one detector focus position, *z* = 0 of the 3D-field distribution at the detector is recorded. Note that we kept 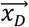 and 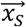 as continuous variables even though in the experiment they assume discretised values, and additional integrations may be needed to account for pixel form factors and scan-induced illumination blur.

Each detector pixel near 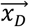 can be interpreted as a small pinhole providing confocal-like sectioning, yielding a transfer function free of a missing cone. This indicates that, with appropriate image processing, optical sectioning can be obtained with this type of ISM data, even though the often-performed pixel reassignment algorithm does not directly provide this.

As seen from Equation (1), the acquired single slice data is best ordered as a 4D-dataset. The first 2-dimensions are the detector pixels while the other 2-dimensions are the scanning positions through the sample. Here, the beam is scanned by shifting the illumination pattern along the x- and y-direction on the projector. In order to create the temporally moving grid of illuminating spots in the sample plane, common ISM setups rely on a large number of carefully aligned optical parts such as lasers, galvanometric scanning mirrors, scanning lenses and mirrors[19]. The aim is to image the scan mirror into the back focal plane of the imaging objective lens, which converts the angular variation into a transversal scanning of the beam. A majority of the parts necessary for the x and y scanning can be omitted by using only one scanning MEMS-based mirror[12]. Here, we rely on two consumer-grade laser video projectors (Figure 2 b) to realize a complexity reduced and low-cost ISM setup. Our used image-processing is based on [20].

**Figure 2 –.**
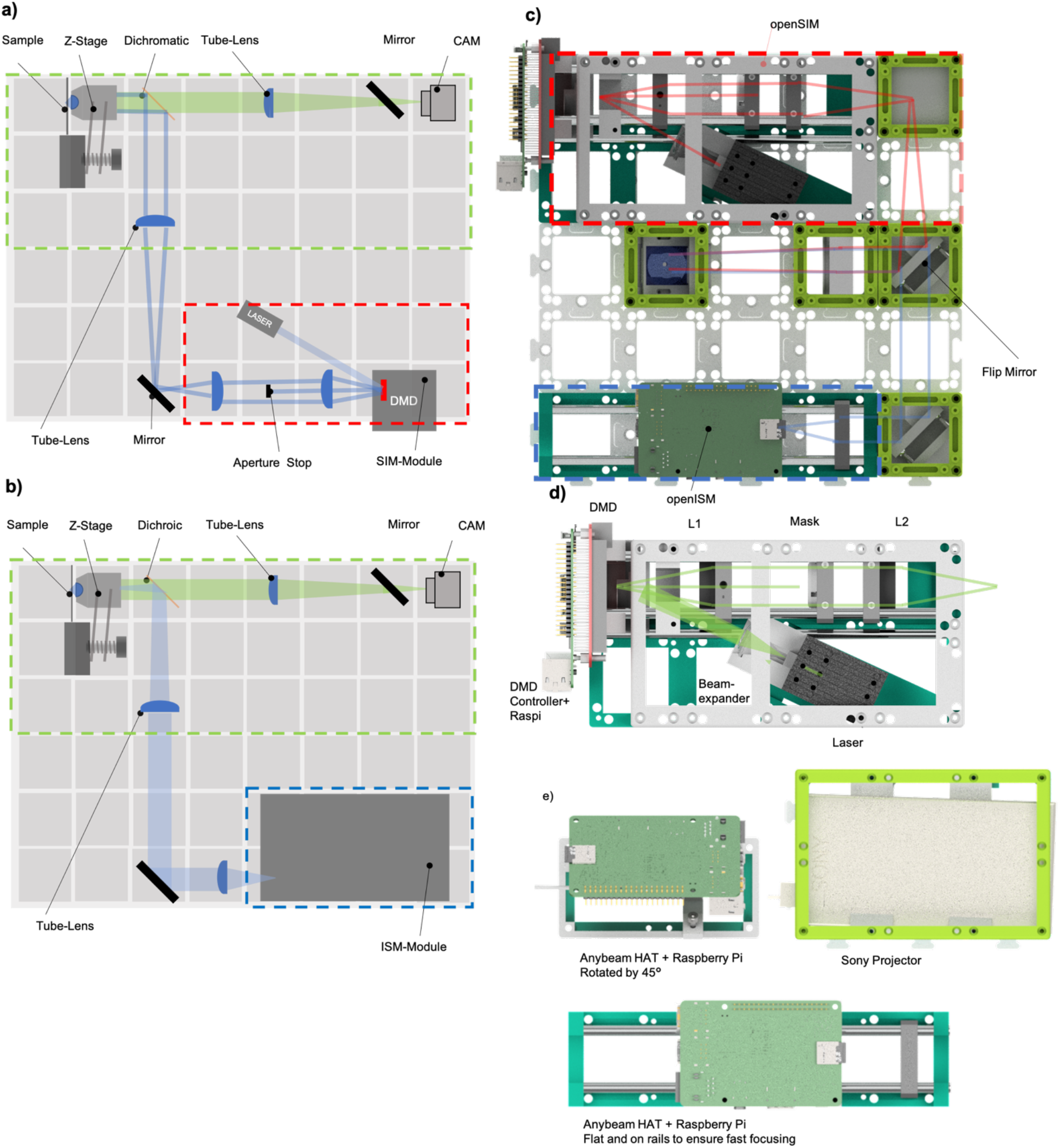
Basic Scheme of the Combined ISM and SIM Setup using UC2: The complexity reduced SIM-setup is a) translated into the UC2 block design and shows b) similarities to the ISM setup. Both designs share the same detection path marked with the green dotted line. The d) openSIM module and both e) openISM modules, equipped with the Anybeam HAT or Sony laser projector, have their own optical UC2 module for c) click-fit adaptation into a setup.

### Structured Illumination Microscopy (SIM)

Structured illumination microscopy (SIM) is another method for optical super-resolution with optical sectioning capabilities^27^, which can improve the lateral resolution up to a factor of two as compared to the standard brightfield imaging^23^. By making use of the Moiré-effect, fine (high-frequency) and potentially not detectable patterns can be down modulated into coarse patterns (low-frequency) by overlaying them with purposely-designed fine (high-frequency) patterns.

In this work we implement 2D-SIM with coherent illumination, where two plane waves are interfered to create the illumination pattern. The m^th^ illumination phase image 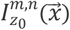 at grating orientation *n* is also described by the top part of equation 1, however, here 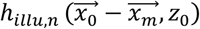 constitutes a sin^2^(·) pattern with grating vector 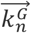 of grating direction *n,* changing phase via various pattern displacements 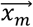. A detailed explanation can be found in [21].

A number of studies have shown that the formerly mechanically shifted grating can be replaced with a spatial light modulator (SLM). This has the advantage that the grating constant, the phase and the grating angle can be freely chosen in order to accommodate multiple wavelengths or different objective lenses. In addition to liquid crystal on silicon (LCoS)-based setups[22], DMD-based devices have gained more attention recently due to their fast switching and displaying speed and their low price[23]. We implemented a DMD-based SIM-module to reduce complexity and footprint (Figure 2 a).

## Results

### UC2 injection moulded cube

Motivated by the implementation of a UC2-based scientific workshop with 150 students and the associated high demand for plastic cubes, we took this as an opportunity to produce the parts using high-volume injection moulding (IM). We consulted a company specialized in rapid prototyping (CNC Speedform AG, Werther, Germany) that helped us translate the 3D printable parts into a design that could be replicated with a minimum number of tools to reduce the cost and complexity of the production process. Production is based on a 250kN injection moulding machine (BOY, 22A, Germany) that uses ABS as the base material. This in combination with a precision-machined aluminium injection mould allows for higher repeatability in mechanical dimensions with lower tolerances compared to 3D-printed parts. We validate this by measuring the external dimensions of 10 randomly selected 3D-printed and IM cubes. The IM cubes have a mean outer dimension of 49.85±0.08 mm, while 3D-printed parts have a significant deviation from the ground truth dimension 49.8 mm with 49.78± 0.17 mm. The same is true for the diagonal of the cube facet, which is supposed to be 48 mm and results in 47.56±0.18 mm for the IM part and 47.05±0.78 mm for the 3D printed part. This confirms that 3D printing exhibits reduced reproducibility compared to IM parts. To mitigate the influence of printer and material types and print settings, we developed a calibration tool^2^, which aims for providing users with the ability to obtain similar printing results across different devices. It helps in the selection of proper printing parameters and correct dimensions for the puzzle piece of the baseplate. This is also a part of the updated module developer kit (MDK^3^) and computer aided design (CAD) documentation. The MDK now relies on the puzzle-shaped baseplate rather than the monolithically printed larger baseplates, which tend to bend after the print cools down.

The advantage of the mass-produced injection moulded cubes lies not only in speeding up the production, which is otherwise around 3,5 hours per 3D printed cube and <1 min for the IM parts, but also in the above-mentioned increased reproducibility. Due to higher precision of the IM cubes, the assembly boils down to pressing the halves of the cube together, closing the insert inside. The inserts can be easily exchanged with almost no effort, unlike the 3D printed cubes where the four screws holding the lid must be disassembled, which quickly degrades the printed material around the threaded holes, hence the whole cube.

To have a better match to the new IM design and to fulfil the requirement of moving the sample rather than the objective lens we developed a new Z-stage, which benefits from the stability of the surrounding IM frame^4^. The motorized (28BYJ-48, China) micrometre-driven (RS Electronics, UK) flexure-bearing offers a 4× reduction and can focus the sample with a step-size of ≈ 900*nm*.

We measured long-term stability of injection moulded vs 3D printed cubes with two identical UC2 Michelson interferometers^5^, one of them using the IM cubes with 3D printed inserts and the other purely with the 3D printed (3DP) components. We expanded a 532 nm single-mode diode laser (5 mW laser pointer, laserlands.net) by a factor of 5 and split it in a 50:50 ratio with a beam splitting cube (optikbaukasten.de, Germany). Both beams are reflected by kinematically mounted (ball magnets on metal plate) cosmetic mirrors and brought to interference on the Raspberry Pi camera CMOS sensor (Raspberry Pi, v2, UK). A customized Python script periodically (i.e. every minute) turns on the laser and records the interference pattern. We visualize the time trace along the centre lines (X/Y) of the 2D stripe pattern in **Fehler! Verweisquelle konnte nicht gefunden werden.** for the IM setup b) and the 3DP setup a).

Both setups exhibited a similar amount of drift over time, especially for a longer time period (~15 hours). The 3DP setup experience a smooth drift over time, whereas the IM components lead to an abrupt change in the contrast (visible in the discontinuities in **Fehler! Verweisquelle konnte nicht gefunden werden.** b), suggesting a sudden motion of one of the inserts. Unfortunately, we were not able to reliably compare the performance for the first hour after alignment, due to the unstable performance of the low-cost lasers in terms of emitted wavelength, phase and intensity. This had a severe but random effect on the contrast of the interference pattern. The lasers were identified as the main cause of the variation in the interference pattern, followed by the 3D-printed laser holder inserts and the kinematic mirror inserts. The IM cubes provide comparable stability to the previous 3D printed system and the variation comes from the other components.

### openISM – Hardware Setup

Both ISM and SIM rely on a temporal variation of the illumination pattern 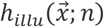 to excite a fluorescent sample 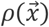, where 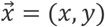 marks the lateral coordinate vector and *n* the number of the displayed pattern. Even though the method to generate the excited field *h*_*illu*_ differs in ISM and SIM, both setups share most of the optical paths, here the back focal plane (BFP) and the whole detection path. We therefore split the setup in three different functional layers, aiming to be used for a single excitation wavelength at a time:

- The **imaging layer**, which hosts the camera, the tube lens and kinematic mirror mounts to adjust the beam path (Figure 2 a/b, green box),
- The **light engine layer**, which includes the laser projector module (openISM) responsible for the creation of the incoherent point-scanning pattern as well as the DMD-based module (openSIM) for the creation of the two-beam interference for SIM and kinematic mirror as well as a dichromatic beam splitter cube to align and filter the light (Figure 2 a, red box, Figure 2 b blue); an inserted flip-mirror cube switches between SIM and ISM mode
- The **sample layer**, which hosts the stationary fixed objective lens as well as a z-stage for precise sample movement.

We choose a MEMS-based laser-scanning projector for the openISM module (Figure 2e). It offers higher contrast than the DMD-based systems, since the laser is only turned on when necessary, thereby reducing the background signal and bleaching. We compare two different consumer-grade video projectors (Sony, CL1A, Tokyo, Japan, 300€; Anybeam HAT, UK, 200€) with a MEMS module from Microvision and Ultimemes respectively. The XY-scanning mirror is magnified into the BFP of the objective lens, which transforms the angular variation into a lateral displacement of the focussed beam in the sample plane. By turning on the laser at distinct scan positions, a lattice-like structure is generated. Slightly overfilling the aperture allows achieving the smallest possible excitation focus (i.e. *h*_*illu*_) in the sample plane. A telescope, consisting of a second achromatic projection lens (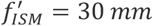 Thorlabs ACS-254 30) and the tube lens *L*_*T*1_ = 165 *mm*, is placed in front of the projector to magnify the circular scan mirror with diameter *D*_*mirror,sony*_ = 1.0 *mm*, *D*_*mirror,anybeam*_ = 1.2 *mm* by a factor of ≈ 5 ×. Due to geometrical constrains of the system, this does not fully meet the requirements to overfill the BFP (i.e. diameter of the entrance pupil, *D*_*EP*_) diameter for both of the two lenses used in our study (Carl Zeiss 63 ×, NA=1.4, *D*_*EP*_ = 6 *mm*; China No Name, 100x, 1.25, *D*_*EP*_=4.5 mm). Additionally, the projection lens (*D*_*Lism*_ = 25 *mm*) cannot fully capture the field of view offered by the projector, which is 43.2° by 24.3° for the Sony and 42° by 24.3° for the AnyBeam.

The commercially available plug-and-play Sony projector is controlled using an HDMI video connection and therefore potentially compresses data connection, whereas the Anybeam offers full control over settings and imaging parameters through their open-source Raspberry Pi API through SSH. Both operate at a framerate of 60 *Hz*, where the Sony offers an uncommon pixel sampling of 1920 × 780 *pixels*^2^ and the Anybeam a resolution of 1280 × 720 *pixels*^2^. Colour modulation is done by mixing three pulse-width modulated (PWM) laser lines[25] *λ*_*R*_ = 635 *nm*, *λ*_*G*_ = 532 *nm*, *λ*_*B*_ = 450 *nm* at a peak power of *P* = 206 *mW* (e.g., white frame) for the Sony and *λ*_*R*_ = 638*nm*, *λ*_*G*_ = 525*nm*, *λ*_*B*_ = 442*nm* at a peak power of *P* = 280*mW* for the AnyBeam, matching the absorption spectra of popular fluorophores such as Alexa Fluor 647, mCherry and GFP. Compared to the Sony projector, which exhibits a freeform optical element in front of the oscillating mirror element to correct for the distorted projected image, the Anybeam HAT light engine maps the rectangular image directly without additional optical elements. We designed two different adapters, which integrate the devices into the UC2 framework, where we had to ensure proper alignment of the optical axis of the Sony projector which is tilted both in X and Y w.r.t. to the device’s housing. One Anybeam module is flipped by 45° to fit into the uc2 dimensions for a 2 × 1 cube (Figure 2e, upper). The other module is designed such that the projector can be moved on 6mm Thorlabs rods to simplify the adjustment (Figure 2e, lower).

The temporally varying pattern displayed by the projector (i.e. grid of point sources) for multi-spot sample-illumination is generated by a customized python script and displayed full screen using the pygame-package from Shinners *et al.*^6^ via the HDMI connection in case of the Sony projector and the Serial Peripheral Interface (SPI) offered by the Raspberry Pi in case of the Anybeam. The exposure time was set to an integer of 1/60s to avoid beating of the laser and pixel readout of the rolling-shutter of the camera (Alvium U1800 158, AlliedVision, Germany). In the case of the Sony projector frames were synchronized by image post-processing, while in the case of the Anybeam a Raspberry Pi GPIO trigger was wired to the camera. Exposure time was typically between 200-500 ms, hence averaging over multiple frames. We choose a unit cell of 10×10 pixels^2^, where one pixel is in the on-state, as a compromise between reasonable signal to noise ratio (SNR), acquisition speed and background signal. The distance between each consecutive illuminating spot position *d*_*pixel*_ = 120 *nm* is close to the Nyquist-sampling, which is important to avoid residual patterning in the result images.

The detection system is based on a classical infinity-corrected microscope design, where the fluorescent signal is filtered using a dichromatic beam splitter (IDEX Health & Science, Semrock FF660-Di02, USA, 285€) and an emission filter (AHF HQ 675/55, Germany, 340€), before an achromatic tube lens (f=156 mm) forms an image on a monochromatic back-illuminated CMOS camera connected to an Nvidia Jetson Nano single-board computer for frame-acquisition and hardware control. To reduce read noise, we actively cooled the CMOS camera with a 12V fan, which otherwise reaches temperatures up to 65°C and results into an increased readnoise level.

### openISM – Imaging Results

For demonstrating the effect of higher lateral and axial resolution of ISM on biological samples, we use SiR Actin labelled[26] HeLa cells and record a ISM imaging data stack. To compare different reconstruction methods to classical widefield fluorescence data (Figure 4e), we use three different reconstruction algorithms:

1. Super confocal imaging[24]: Here, the stack is processed by applying *max*_*z*_(*I*_*stack*_(*z*))+*min*_*z*_(*I*_*stack*_(*z*))−2\**mean*_*z*_(*I*_*stack*_(*z*)). It exploits the fact that a sample position in ISM is most of the time not illuminated when in focus in this sparse illumination scheme. This successfully reduces background contributions and offers a fast and simple way to obtain good sectioning (Figure 4f).
2. Classical ISM photon reassignment: We base our algorithm on work by McGregor *et al.* [20]. Here, we first register individual illumination positions (i.e. peaks, Figure 4a) in each frame, before a grating is fit into this pattern to recover undetected illumination spots at the gratings crossings and to allow for distortions (e.g. field curvature) originating from the freeform optics or improper alignment (Figure 4c). A snippet of 11 × 11 *pixels*^2^ around each detected peak position is cut-out, multiplied with a a 2D Gaussian to simulate a “macro-pinhole” [20], [27] for better axial sectioning, summed and reassigned on a two times super sampled canvas (Figure 4c). Reassigning the information for all illumination spots in each frame produces a super-resolved image (Figure 4 d). The open-source customized algorithm is available in our Github repository^7^.
3. SRRF: The previously published algorithm by Gustafsson *et al.* [28] offers a simple way to create super-resolved images from image stacks at varying emission signal. For a full mathematical derivation of intensity fluctuation-based super-resolution imaging, the readers is referred to [28],[29].

**Figure 3 –.**
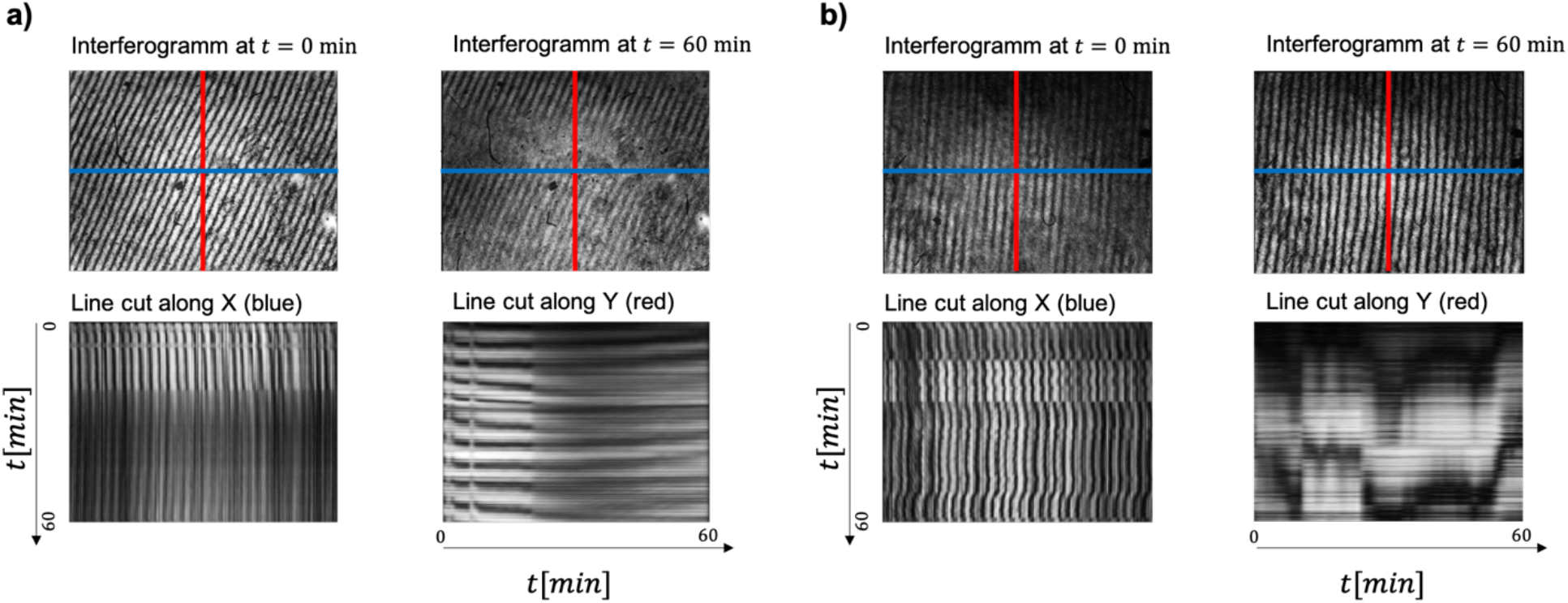
Observing interference of a 3D printed and injection moulded Michelson interferometer – a) shows the temporal variation of the two-dimensional interference pattern of a Michelson interferometer based on 3D printed components equipped with ball magnets and cylindrical screws. In the line cut along X (blue line) and Y (red line), it is visible, that the pattern remains stationary over the timespan of 60 minutes. Only small drift is visible. b) the same experiment was performed with injection moulded cubes instead simultaneously. In the line plots, it is visible, that the pattern is temporally moving at multiple time points which could be due to abrupt motion in the room. Yet in a), the contrast of the pattern stays almost constant over the entire duration of the experiment.

**Figure 4 –.**
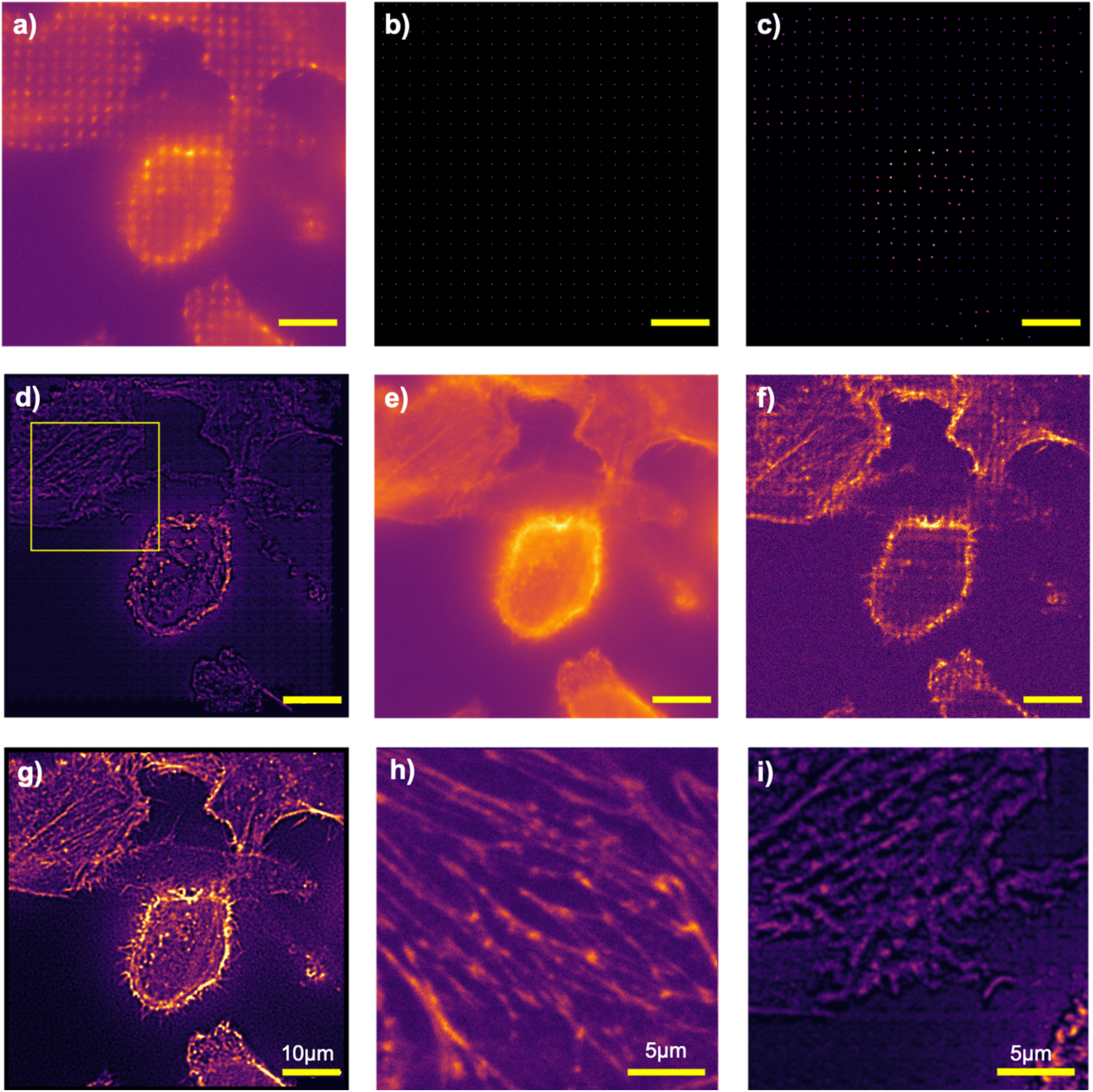
Comparing Imaging Results using different reconstruction methods from openISM (Sony) and an ISM setup based on high-quality components: In a) RAW openISM frame where the illumination spots are clearly visible. b) indicates the fitted peak positions as a grid of points. Around each point a portion is cut-out, weighted using a 2D Gaussian and placed on a two-fold super sampled grid c). d) represents the sum of all filtered intensities across the temporal stack, which is called the ISM reconstruction. Compared to the brightfield image in e), the ISM reconstruction has a reduced background and offers better optical sectioning. The actin fibres in d) are much more visible. f) shows the super confocal result as described in [24], which offers good optical sectioning at a very much reduced computational effort as compared to the SRRF method applied to the same dataset in g). SRRF, similar to classical ISM, can improve lateral and axial resolution, but requires high computational power and tends to produce fibrous results. h) Comparing the imaging results of the same sample between openISM in i) (zoomed ROI from g) and a home-made ISM setup equipped with an sCMOS camera and a scanned laser assembly, shows similar optical performance in terms of resolution and sectioning.

All reconstruction algorithms (1., 2. and 3.) successfully suppress the background signal, which masks smaller object features such as the actin filaments in the homogeneously illuminated images (Figure 4e). Method 1 has the advantage of being computed nearly instantly without any parameters required by the user. This enables the use on devices equipped with low computational power. Method 2 on the other hand serves as a customized algorithm for non-perfect data with respect to the illumination pattern and can increase optical resolution practically up to a factor of √2. Small object details such as the actin filaments are clearly recovered. However, the current implementation of the algorithm is not optimized for fast computation; therefore, a reconstruction of a typical stack of 512×512×100 images takes around 2 minutes on a standard Laptop. Artifacts such as the visible grid-like pattern superimposed to the reconstructed object are caused by improper selection of imaging parameters, e.g., due to unknown interpolation of the laser projector through HDMI, or due to weak contrast of the illumination pattern. They are clearly visible in both, the ISM reconstruction (Figure 4d) and even more dominant in the super confocal image (Figure 4f). Method 3 has proven to be a complexity-reduced super-resolution method specifically powerful for fibrous samples[29]. It directly runs inside Fiji[30] and produces good results with default parameters. We apply the algorithm to the same image stack as the other methods and achieve increased optical sectioning. The overall result looks similar to the total internal reflection fluorescence (TIRF) images (Figure 4g) and does not require any estimation of the illumination pattern, which is a major error source for the ISM reconstruction algorithm.

In addition, we compare the ISM reconstructions of HeLa cells labelled with Alexa Fluor ^®^ 647 phalloidin, between the low-cost setup (Sony) and an existing home-built ISM system based on a modified commercial confocal microscope detecting at the pinhole plane via a very small region of interest of an emCCD at each scan position is slow (180s for a 512×512 image). Here, the 640 nm laser diode laser (LDH-D-C-640; Picoquant, Germany) is scanned in X/Y in the sample plane with a galvo/resonant scanning mirror pair. The de-scanned signal is recorded on a 12 × 12 *pixel*^2^ ROI of a sCMOS camera (Andor Neo, UK). The ISM setup based on high-quality components scans the ROI with 512×512 scan positions with a sampling of ~43 nm, whilst the image acquisition frame rate is limited by the exposure time and data transfer rate of the sCMOS camera and takes about 3 minutes. In contrast, the homemade ISM takes about 1 minute to acquire the 512×512×100 images, while the framerate is only limited by the unit-cell size and not the ROI as it is the case for the point-scanning ISM. With a smaller unit cell size, i.e., the distance between two on-pixels, the acquisition time can be further reduced. Since no additional alignment of the laser w.r.t. the scanning mirror needs to be performed, the UC2 system is much easier to assemble and very compact. Compared to the ISM result Figure 4h), the openISM offers a much larger ROI, but requires additional post-processing steps, such as the registration of the illuminating spots. For sparse samples, this can lead in an incorrect reconstruction result, impossible in the classical ISM setup. The ISM result does not show any periodic artefact in the reconstruction, which is the case for the super confocal result (Figure 4f) and the ISM reconstruction (Figure 4d/i) of the UC2 setup.

During the experiments with the AnyBeam projector, we observed an unpredictable periodic jitter of the projected-generated pattern^8^. We believe this is due to a common problem of raster scanning schemes of MEMS and needs to be investigated more in detail in the future. The accompanying motion blur of the single point foci over several frames causes a severe drop in fluorescent signal, hence not producing high enough SNR for later reconstruction. A firmware upgrade of the device could likely solve this issue in the future.

### openSIM – Hardware Setup

We base the openSIM module on a monolithically printed 5×2 base (Figure 2c), which hosts a low-cost single-mode diode laser (*λ* = 635/637 *nm*, Laserlands.com) and a Raspberry Pi driven DMD module (Texas Instruments, DLP2000EVM, Texas, USA) to generate a sinusoidal pattern for super-resolution microscopy. It directly fits to the UC2 grid-system allowing for a simple integration into existing setups by interfacing with adjacent Fourier- or image planes.

The laser-beam, with a diameter of 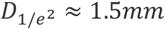, is expanded by a factor of ≈ 6 × with a homebuilt telescope consisting of an iPhone 5 objective lens (*f*’ ≈ 3 *mm*) and a plan-convex lens (*f*’ ≈ 20 *mm*) to overfill the active area of the DMD and to offer a relatively homogenous illumination using the line profile of the laser diode. This assembly can be aligned independently before adding it to the SIM module via a 3-point magnetic bearing mechanism, which helps to adjust the optical axis of the collimated beam being parallel to the base plate and perpendicular to the DMD-plane. Flipping the DMD by 45° around the optical axis and introducing an angle of 25° between the plane of the DMD and the illuminating laser-beam is required to operate the DMD at its design-configuration and maximizes the diffraction efficiency into the ±1^*st*^-order[31],[23]. The DMD plane is oriented perpendicular to the outgoing optical axis and the mirrors in the “ON”-state are thus oriented at 12.5° w.r.t. to the illuminating beam and also the outgoing optical axis. Currently no further wavelength-dependent optimization of the angle was performed. The display is re-imaged by a second telescope consisting of two identical achromatic lenses (*f*′ = 50 *mm*, Thorlabs ACS-254 50). A movable magnetically held Fourier-filter, necessary to block the zeroth order, is placed on a ferro-magnetic plate located in the intermediate focal plane. Higher orders were blocked with the disk-shaped aperture, while a small drop of tin added to a 0.1 mm copper wire successfully blocked the zeroth order. The magnetic attachment allowed for a simple adjustment of the Fourier-filter in x/y to suppress the central order of the illumination light in the pupil plane. The remaining focussed diffraction orders are then reimaged into the pupil position of the objective lens as described earlier for the ISM setup (Figure 2b).

The DMD is driven by a Raspberry Pi using a customized printed circuit board (PCB) and displayed the various patterns (e.g., phase, rotation) generated using a customized python script and the pygame library for display. We here limited ourselves to a maximum of two rotational angles (i.e. 0°, 90°), as the setup was meant for demonstrating the effect not optimizing it to its full potential. Furthermore, no additional elements for polarization control (e.g., linear, circular or pizza polarizer) were used to keep the system’s price and complexity low. Due to the short focal lengths of the 4f system after the DMD (f=50 mm), the ±1^*st*^-orders of the SIM-pattern are close to the central order, thereby making alignment of the Fourier mask complicated and could lead to potential blocking of Fourier-Frequencies for other rotational angles. The grating constant was determined by imaging the two-beam interference on a negative USAF target (Thorlabs, R3L1S4N, USA) which results in a grating constant of *d*_*SIM*_ = 495 *nm*, roughly twice the Abbe diffraction limit *d*_*Abbe*_ = *λ*_*em*_/(2*NA*) = 665*nm*/(2 · 1.4) = 237 *nm*.

During our study, we tested different high- and low-cost objective lenses, where low-cost non-branded lenses (e.g., 100×, 1.25NA, China) offer good quality in the centre of the ROI but suffer from significant field curvature at the edges of the FOV and a reduced transmission. The reason is found in the non-coated lenses, which are not explicitly specified for fluorescent imaging, therefore exhibit a lower transmission for the emitted signal by the fluorophore, resulting in faster photo bleaching of the sample since the excitation intensity needs to be maximized to get sufficient signal for the camera.

A comprehensive set of instructions on how to assemble and align the system, as well as all 3D-printable files and code can be found In the UC2 hardware repository[32], [33].

### openSIM – Imaging Results

For demonstrating the resolution enhancement using the openSIM module, we use Alexa Fluor ^®^ 647 labelled HeLa cells and compare reconstructed results with those from a commercially available SIM system (Zeiss, Elyra S.1, Germany). We acquire a stack of 4 images at varying phase, where the grating constant was chosen to be 495 nm using the Zeiss 63 ×, NA=1.4 objective lens to display sectioning capabilities and resolution enhancement of our implementation of only one rotational angle of 2D-SIM.

The data-sets were reconstructed with a custom MATLAB toolbox based on DipImage [34] (available upon request) which automatically detects experimental parameters such as grating constant, phase and rotational angle. Alternatively the use of the open-source Fiji Plugin fairSIM [11] is possible (see Supporting Figure 1).

The comparison of a single raw frame (**Figure 5a**) with the widefield (Figure 5b) and SIM-reconstruction (**Figure 5c**) displays clear recovery of fine details of the Actin network along one direction (**Figure 5c**). Compared to results of the commercial device in Figure 5d), which is almost an order of magnitude more expensive, the openSIM reconstruction lacks a uniform gain of optical resolution, since we use only one grating orientation. Additionally, since the active area of the DMD is relatively small and uses only one half of the active sensor area of the camera, the field of view is much smaller compared to the Zeiss Elyra. Similarly, to the comparison of the openISM reconstructions, we compare reconstructions of SIM data (3 phases, 1 rotational angle) using the original SIM reconstruction algorithm (**Figure 5**e) and the SRRF algorithm directly applied to the SIM raw data (Figure 5f). SRRF has shown to work well on fibrous samples without changing the default parameters and produces a higher uniformity of the recovered optical resolution and intensity distribution.

**Figure 5.**
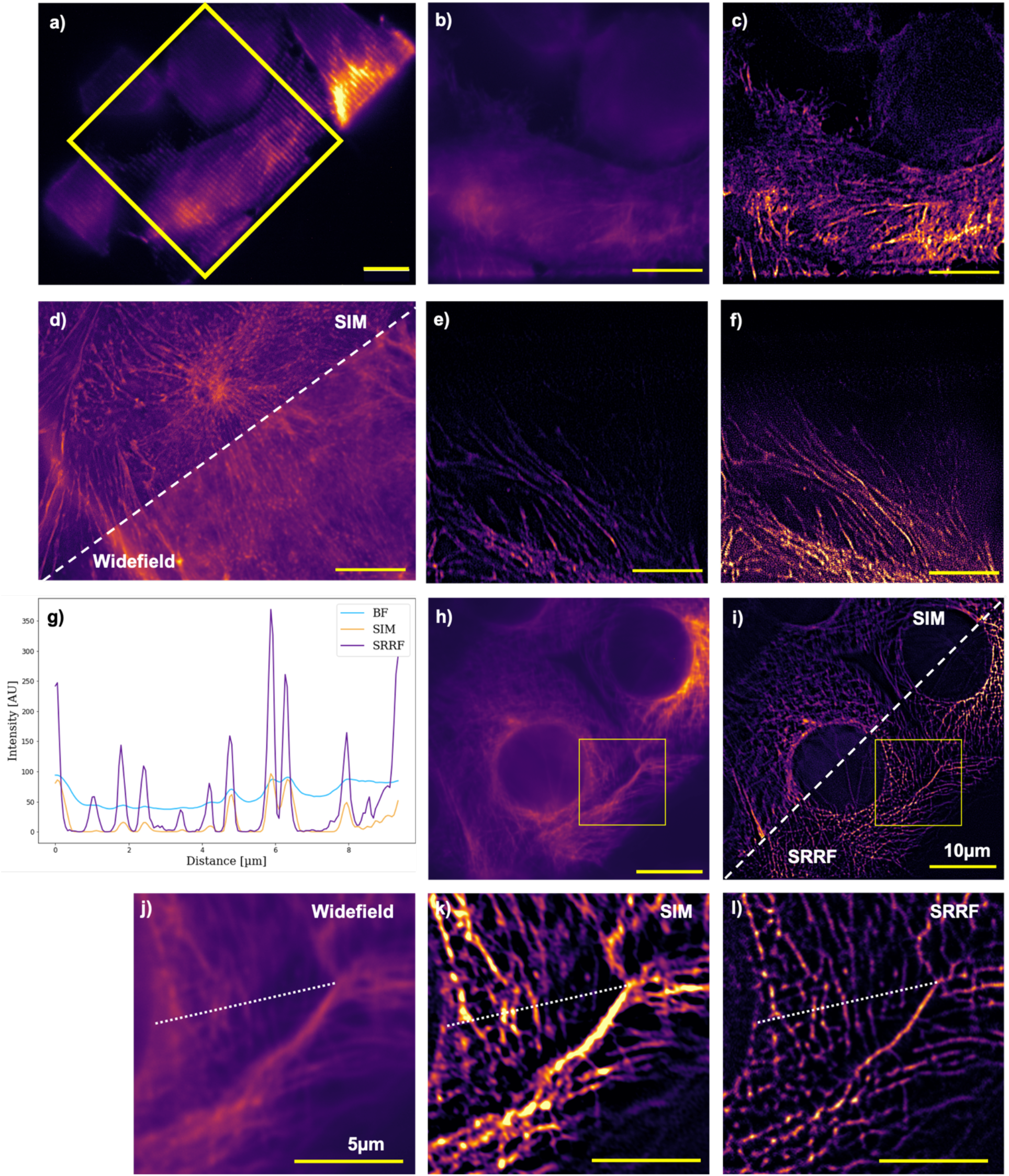
– Comparing Imaging Results using different reconstruction methods from openSIM and a commercial SIM system: a) RAW SIM frame from the UC2 openSIM system of AF647 labelled HeLa actin network is acquired high-quality with an objective lens (Plan-Apochromat 63 ×, *NA* = 1.4, Carl Zeiss, Germany) at λ_exc_ = 635nm. The illumination grating is aimed to produce a 1.5 × resolution improvement. b) The 45° tilted widefield (=sum) image and c) the SIM reconstruction (Wiener-filter parameter[22] |k|^0.7^, 3 phases,1 rotational angle) using the openSIM module. In the latter actin fibres are clearly recovered and out-of-focus background is suppressed. d) A comparison between widefield (right) and SIM reconstruction (left) of the same sample, but different ROI using a commercially available Zeiss Elyra S.1 with the same objective lens. e) SIM and f) SRRF reconstruction of openSIM data of different sample of same type. h) Widefield image of microtubule labelled (AF647) HeLa cells and i) the SIM (left) and SRRF (right) reconstruction of the image stack with four phases and two rotational angles (0°,90°). j-l) Zoomed in region of interests from the yellow box in h-i) to compare the visible details in widefield, SIM and SRRF. g) The line plot along the white dashed line in j-l) suggests a line-profile at full width half maximum (FWHM) of d_FWHM,widefield_ = 289nm, d_FWHM,SIM_ = 140nm, d_FWHM,SRRF_ = 136nm.

Careful alignment of the Fourier mask enables the acquisition of multiple rotational angles (i.e. 0°, 90°) and offers higher uniformity in optical resolution as demonstrated in **Figure 5 h-l**), where we used microtubule labelled HeLa cells (α-tubulin/α-mouse AF647, ThermoFisher, A-21235, USA). In contrast to actin labelled samples, the new dye offered an increased and more stable signal, leading to a reduced exposure time (*t*_*exp*_ = 350*ms*) and significantly reduced photobleaching due to lower laser power. Comparing the widefield (**Figure 5h**) with the SIM- and SRRF reconstructions (Figure 5i) on the image stack (4 phases, 2 rotational angles), the reconstructed images show a much better uniformity in resolution improvement compared to one rotational angle-only SIM results in Figure 5c/e). The resolution, measured as the full width half maximum (FWHM) at multiple location of the microtubule network leads to a resolution of *d*_**FWHM**,*widefield*_ = 289*nm*, *d*_**FWHM**,*SIM*_ = 160 *nm* and *d*_**FWHM**,*SRRF*_ = 156*nm* slightly surpassing the aimed resolution improvement of 1.5 ×. A further advantage can be seen in the suppressed background signal of the reconstructed data, leading to a better optical sectioning.

An imaging result using the 100× low-cost objective lens can be found in the Supporting Figure 1. The result of the SRRF algorithm, directly applied to the 4 SIM raw images, clearly recovers the microtubule structure of the cell samples, weather the SIM reconstruction using fairSIM suggests a strongly varying grating contrast across the sample, hence leading to reconstruction artefacts and a failed reconstruction. Decreasing the grating constant, hence lowering the gain in optical resolution helps to increase grating constant and can lead to an increased optical sectioning using this low-cost setup. With this, the SIM and SRRF reconstruction can clearly resolve the microtubules structure by successfully blocking the background signal.

## Discussion

We presented a mechanical upgrade (UC2 v3) to the recently introduced modular optical toolbox UC2, which improves long-term stability by introducing injection moulded (IM) parts with much lower manufacturing tolerances thereby increasing reproducibility of optical setups. From our practical experience in the optics lab, mechanical stability can keep up with metallic cage systems, only that individual modules, such as lenses, can be easily replaced by swapping a new one in. We assured to keep compatibility between the IM parts and the 3D-printable parts by providing a 3D-printed version of the puzzle piece-like assembly of the baseplate and the additional pin-based mounting mechanism. We updated our modular developer kit (MDK v3) and introduced a calibration tool to make the form locking mechanism available to the 3D-printed cubes. We used the same 3D-printer (PRUSA i3 MK3S, Czech Republic) for all 3D-printed elements, but the used PLA-filaments seem to differ slightly between the vendors and therefore the parameters had to be adjusted per batch. Even after calibration the tolerances of the 3D-printed parts where still higher than the IM parts. We observed comparable drift of the IM based and 3D printed-only setup. More reliable external components and potentially also injection moulded cube inserts would be necessary to judge the performance quantitatively.

During experiments we noticed that the increased mechanical stability of the IM parts eventually reduced the flexibility in exchanging parts force-free. Some parts held together so strongly that an increased amount of force was needed to loosen them, thus posing the risk of misaligning other parts of the optical assembly. This is especially true for the dichromatic beam splitter surrounded by cubes and baseplates on all sides. Switching to another wavelength requires a dis- and reassembly of many parts, followed by an additional alignment step. Unlike with the 3D printed cubes, there is no risk of accidental misalignment or misplacement of a cube, thanks to the firm mechanism that attaches the cubes to the baseplates, making it very easy to transport the system with only little misalignment as observed with the IM-based Michelson interferometer, which showed interference even after putting it in a box, transporting that to another lab. The current micrometre-driven flexure bearing z-stage was designed to be integrated into existing UC2 cubes, hence limiting the dimensions of samples compatible with the system. This must be accounted for when preparing samples. A 35 mm Petri-dish, for example, will not easily fit into the system. Wavelength-depend design (the order selection apertures) and alignment differences can easily be compensated by providing multiple openSIM modules. Each module can host a single laser line (e.g. available low-cost diode lasers: 450nm, 488nm, 520nm, 532nm, 635nm) each featuring an individually optimized relative angle between the incoming light and its individual stationary DMD to account for individual wavelength optimized blaze conditions.

The quality of the resulting SIM reconstructions depends on the quality of the created excitation pattern in terms of contrast and homogeneity. Since the 3D printed module did not permit large variations in the optical alignment, the blaze condition was not yet perfectly matched, which caused one of the first order beam only has 50% of the intensity of the other first order degrading the grating contrast. Also imaging artefacts, like inhomogeneity of the emitted signal due to a slightly misaligned optical axis degrade the reconstructed result. In principle, the use of cheaper chromatic filters (e.g. Comar 740 IY 116, UK, 30€) is possible. However, in this case the system suffers from a considerably lower SNR, as the transmission rate (50% at the *λ*_*em*_ = 660*nm*), is much lower. However, it is worth further investigation into low-cost alternatives as filters are a significant cost factor for this system. The two rotation angles used for the SIM pattern at 50% of the frequency limit (for optimal sectioning) limit the resolution enhancement to a theoretical factor of ≈ 1.42 × along the diagonal direction compared to 1.5 × along the pattern orientation directions. However, this follows in a considerably simplified adjustment, since the Fourier aperture otherwise quickly runs the risk of accidentally blocking one of the necessary orders. The reason is found in the selection of lenses with short focal length for the 4f system required to apply a Fourier filter. Due to the limited separation of the diffraction orders, adjusting the position of the Fourier mask is becoming difficult, since blocking parts of one of the used orders becomes very likely. Lenses with a longer focal length could help, but they would require a larger setup.

With the newly introduced increased mechanical stability, we demonstrated that optical super-resolution methods like ISM and SIM can be made widely available with off-the-shelf components. Therefore, existing optical setups were translated into the compact block-based system. We found a low-cost alternative for every component in the original setup leading to an overall cost of around 2000 € (see Table 1) including all optics, electronics and computers to acquire and process the images.

Nevertheless, the low cost comes at a price of reduced flexibility when it comes to adjusting experimental parameters. The here presented consumer-grade laser projector (Sony) was a limiting factor in the openISM setup. Its black box-like behaviour in terms of pixel-addressing, interpolation-scheme and the optical properties of the freeform lens in front of the laser scanner complicate integration in the optical system and experimentation. The recently introduced Anybeam HAT solves this problem by granting full access to the hardware. Nevertheless, an in-detail characterization is still pending. During our experiments we faced an unpredictable jitter of the pattern, which needs further investigation. Also, the selection of low-cost objective lenses not optimized for fluorescent imaging poses the problem of low transmission of fluorescence signal, leading to an increased bleaching rate, since the laser intensity needs to be adjusted to get sufficient signal for the camera. The current setup does not include a simple mechanism to correct a potential mismatch in the blazed angle, therefore a reduced contrast in the SIM pattern was observed, also evident from the mismatch in intensity in the different diffraction orders used for the interference.

## Conclusion

The newly introduced injection moulded cubes and baseplates allowed us to build a highly compact and stable combined super-resolution optical setup. With openSIM and openISM we realized two new contributions to the UC2 ecosystem and demonstrated how more sophisticated optical setups can be made widely available. In our study we decided to balance resolution gain against optical sectioning to achieve a stable and impressive imaging performance. The selection of low-cost components still limits the use of this setup for productive use and a realignment of the optical path is needed frequently. With this module we are not yet aiming for front-end optical resolution, but rather for the use in educational areas and for rapid prototyping where data generation (e.g. for machine-learning) or development of image processing algorithms is of interest. Having access to all adjusting screws and experimentation parameters makes this a valuable tool for education, teaching and outreach, since the financial impact of a damaged part is not as severe compared to its commercial counterpart.

We believe that this contribution to the UC2-ecosystem displays the possibilities of the toolbox and marks the next step towards front-end research with frugal methods.

## Supplementary

### Sample preparation

#### HeLa cell preparation

Cells were cultured in Dulbecco's Modified Eagle Medium (DMEM, Thermofisher, USA) supplemented with 10% Fetal bovine serum (FBS) and 1% Penicillin/Streptomycin Solution (P/S) and maintained in 37 ℃/5 % CO2 incubator. Cells were harvested at 90% confluency. The culture dish was first washed with 1 ml Dulbecco’s phosphate-buffered saline (DPBS, Thermofisher, USA) twice and incubated in 1 ml TrypLE (Thermo Fisher, 12604013, USA) at 37℃/5% CO2 for 5 min. Afterwards, 8 ml fresh culture medium was added to the dish to deactivate the TrypLE. 1 ml of the cell suspension was transferred into a new dish with 9 ml medium and the rest of the cells were transferred into a glass bottom chambered well plate (μ-slide, ibidi, Germany) which was prepared for fluorescent labelling. The culture medium was changed every 4 days.

#### HeLa + SIR

Silicone rhodamine (SiR) is a deep red fluorophore for super resolution imaging [29]. The HeLa cells were stained in culture medium with 1 mM SiR at 37℃/5% CO2 for 1 hour. Later the cells were washed with 1 ml DPBS twice and fixed with 4% Paraformaldehyde (PFA) diluted in PBS for 15 min. The sample was washed three times with 40 μL PBS for 5 min. The fixed cells were stored and imaged in PBS solution.

#### HeLa + AF647 phalloidin

Alexa Fluor 647 is a fluorescent dye with the similar excitation and emission spectra as SiR. To prepare the samples with AF647 phalloidin, the culture medium of HeLa cells was first removed and washed with 0.4% Glutaraldehyde (GA) and 0.25% Triton X-100 solved in PBS for 90 second. The sample was following fixed with 3% GA in PBS for 15 min. Then the cells were washed with pure PBS for 3 times to clearly remove the GA and blocked with 0.3 M glycine in PBS for 10 min. The sample is stored and imaged in PBS solution.

#### HeLa + Anti-Mouse IgG antibody AF647

Anti-Mouse IgG secondary antibody AF647 was used to label the microtubule of the HeLa cells. The HeLa cells were first washed with pure PBS for three times, then followed with 0.4% GA and 0.25% Triton X-100 solved in PBS for 90 second. The sample was fixed with 4% PFA for 15 min in room temperature and washed with PBS three times to totally remove the PFA. The nonspecific binding sites of the cells were blocked with 3% Bovine Serum Albumin (BSA) for 30 min and again washed with PBS for three times. The sample was labelled with 1:200 primary IgG antibody (Thermofisher, invitrogen, USA) at room temperature for 1 hour, followed by an additional washing step with PBS and then labelled with 1:250 anti-Mouse IgG secondary antibody for 1 hour. The sample is stored and imaged in PBS solution.

## Additional Information

**Information on the following should be included wherever relevant.**

## Acknowledgments

We thank Alice Sandmeyer and Marcel Müller for fruitful discussions about DMD-based SIM setups and image processing. We thank Michael Klein from CNC Speedform AG for his generous support when translating 3D printing files to injection moulding designs. We thank Monalisa Goswami for imaging the HeLa cells using the homebuilt ISM setup. We want to thank Patrick Then for helpful comments on sample preparation.

## Data Accessibility

ISM-Code and Imaging data: https://github.com/bionanoimaging/UC2_ISM_CODE

UC2 openSIM module: https://github.com/bionanoimaging/UC2-GIT/tree/master/APPLICATIONS/APP_openSIM

UC2 openISM module: https://github.com/bionanoimaging/UC2-GIT/tree/master/APPLICATIONS/APP_openISM

Experimental data: https://zenodo.org/record/4041339 (DOI will be updated once peer-reviewed)

SIM-Code: Available upon request

## Authors’ Contributions

Theoretical model, performing experiments, hardware design, assembly, instrument validation and documentation HW, BD, BM. Software development, original concept and project management, RH, BD. Sample preparation and imaging: HW, BD. Writing of manuscript: HW, RL, BM, RH, BD. Funding acquisition: BD.

## Supporting Table 1

**Table 1:**
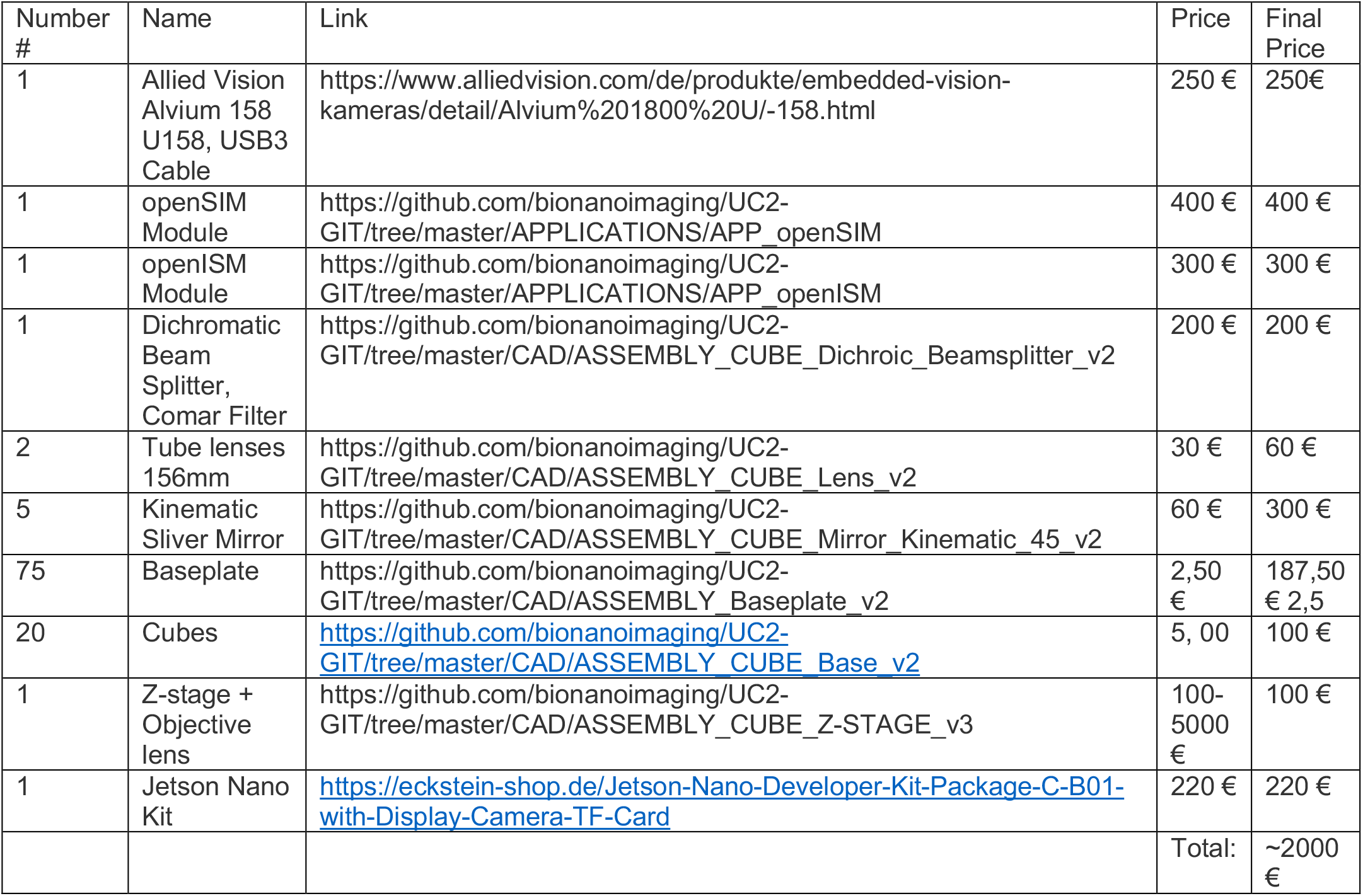
A coarse summary of the modules and components used in this study. A more detailed list can be found in the online repository of the UC2 system.

## Supporting Figure 1

**Supporting Figure 1 –.**
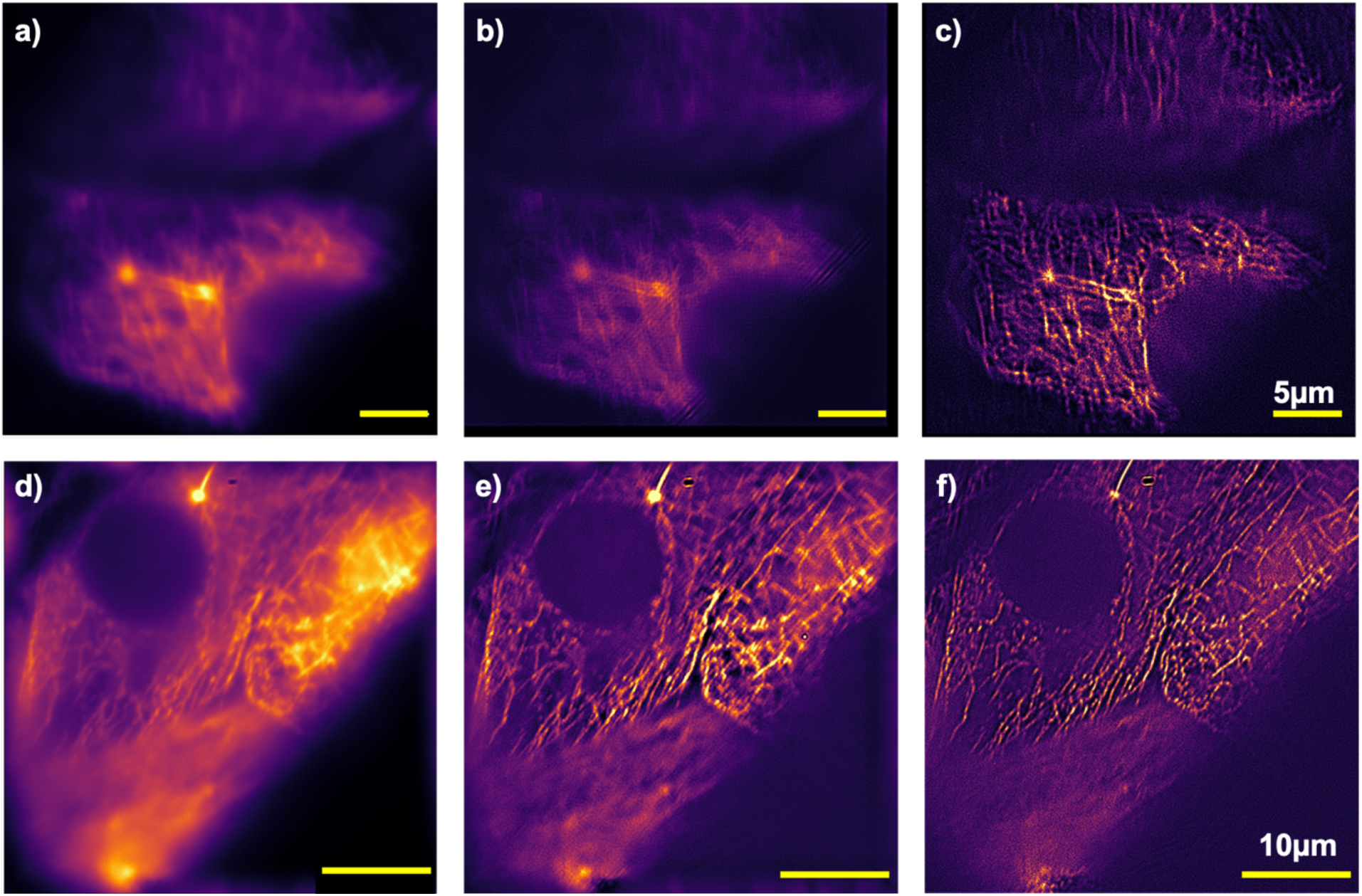
Imaging and reconstruction results using the low-cost 100× objective lens: a) The filaments in the widefield image acquired with the low-cost 100×, 1.25NA objective lens (80€) show a significantly increased blur and aberration compared to the top-shelf 63×, 1.4NA lens used previously. b) The fairSIM reconstruction has problems to estimate the parameters correctly. Reconstruction artifacts, such as the superposed grating is clearly visible, whereas the SRRF reconstruction applied to the same stack of 8 SIM images (4 phases, 2 rotational angles) can clearly recover the micro tubulin structure. Increasing the grating constant to d_SIM_ = 619 nm and the number of phases to 5, which leads to reduced enhancement of optical resolution, the diffraction orders much better matches the BFP of the low-cost objective, leading to better contrast of the grating in the sample plane. The widefield image of a different region of the micro tubulin sample in d) can hardly resolve the filament structure due to increased background level. The MATLAB-based SIM reconstruction in e.) and the SRRF reconstruction in f) can enhance optical sectioning and lead to a recovery of theses details.

## Footnotes

1 https://github.com/bionanoimaging/UC2-GIT

2 https://github.com/bionanoimaging/UC2-GIT/tree/master/CAD/ASSEMBLY_Baseplate

3 https://github.com/bionanoimaging/UC2-GIT/tree/master/MDK

4 https://github.com/bionanoimaging/UC2-GIT/tree/master/CAD/ASSEMBLY_CUBE_Z-STAGE_v3

5 https://github.com/bionanoimaging/UC2-GIT/tree/master/APPLICATIONS/APP_Michelson_Interferometer

6 www.pygame.com

7 https://github.com/bionanoimaging/UC2_ISM_CODE

8 https://github.com/NebraLtd/AnyBeam/issues/14

